# Cortico-hippocampal representational similarity contributes to learning of multiple motor sequences

**DOI:** 10.1101/2023.08.28.555219

**Authors:** Seojin Yoon, Sungshin Kim

## Abstract

Cortico-hippocampal interaction is integral to memory consolidation during sleep or wakeful rest. However, the extent of its engagement in non-declarative motor memory during skill learning remains largely undefined. Specifically, it remains unclear how neural representations in cortical and hippocampal regions are related to learning multiple motor sequences. Here, we conducted an fMRI experiment in which participants learned four distinct motor sequences in a pseudo-random order with interleaved rest periods. We found pattern separation in the sensorimotor regions was significant, particularly in the anterior motor cortex, related to early learning performance. However, there was no pattern separation in the hippocampus during the rest periods. Intriguingly, the representational similarity between the motor cortex during task periods and the hippocampus during rest periods was highly correlated with overall learning performance. These findings suggest a new role of hippocampal replay in motor sequence learning, that is, learning a more abstract relational structure of sequences instead of representing sequences per se.

## Introduction

The motor memory consolidation process involves the transformation of initial fragile memory into more stable long-term memory, which occurs within a few hours after the completion of training ^1^. Previous neuroimaging studies have suggested sensorimotor regions such as the primary motor cortex^2,3^, striatum, and cerebellum^4^ as the neuronal substrates of the consolidation. Furthermore, noninvasive brain stimulation applied to the primary motor cortex (M1) can either disrupt5 or enhance^6,7^ consolidation, further clarifying its causal role in memory consolidation. The striatum and cerebellum also play roles in overnight memory consolidation, with the former associated with motor sequence learning and the latter linked to motor adaptation, respectively^4^.

These studies focused on memory consolidation over long-term timescales within the sensorimotor regions during sleep. However, other neuroimaging studies have found evidence that the hippocampus, previously thought to be independent of motor memories, is involved in sleep-dependent consolidation^8,9^. Moreover, more recent studies reported the role of the hippocampus in a faster form of consolidation during awake rest periods interleaved with practice^10-12^. The rapid consolidation may contribute to short-term performance improvements during the rest periods, known as micro-offline gain. Neural replay or reactivation during the rest periods could be a mechanism underlying the rapid consolidation of motor memories^12,13^. The reactivation occurs not only in the hippocampus but in the cortical regions related to learned information^14-16^.

Indeed, the cortical-hippocampal interaction is central to the complementary learning system (CLS), integrating new information processed by the hippocampus into the long-term stable memories in the cortical region^17^. Human EEG-fMRI studies^9^ and a recent animal study^18^ suggested functional coupling between cortical regions and the hippocampus as the mechanism of motor memory consolidation during sleep. However, these studies focus on the role of cortico-hippocampal interaction in sleep-dependent memory consolidation at a long-term scale. Thus, it remains unclear how the interaction hypothesized in the CLS theory contributes to rapid motor memory consolidation during wakeful rest periods.

Here, we designed a human fMRI experiment where participants practiced multiple motor sequences simultaneously with interleaved rest periods. We analyzed how representational patterns of the sequences in the sensorimotor regions during active task periods mirrored those in the hippocampus during the wakeful rest periods. Further, we assessed how these activity patterns in the sensorimotor regions and the hippocampus correlated with individual learning performance. We discovered that both the cortico-hippocampal representational similarity and pattern separation of multiple motor sequences in the anterior motor cortex were highly predictive of learning performance. These findings indicate that the cortico-hippocampal interaction is integral to the rapid consolidation of motor memory, contributing to improvements in learning performance.

## Results

### Successful learning of four motor sequences

In the MRI scanner, 25 participants performed a well-established motor sequence task^10-12^, which consisted of a series of trials where they had to alternate between 10 s active task periods and following 20 rest periods. During the active task periods, we instructed them to tap a sequence of eight items multiple times using their non-dominant left hand while the items were displayed on the screen (Figure 1A). Four different sequences were presented in a pseudo-random order, i.e., each was presented once in four consecutive sequences. A block of four sequences was repeated six times in each of the four fMRI runs, resulting in a total of 96 trials, taking approximately 50 minutes, including a break between fMRI runs.

**Figure 1.**
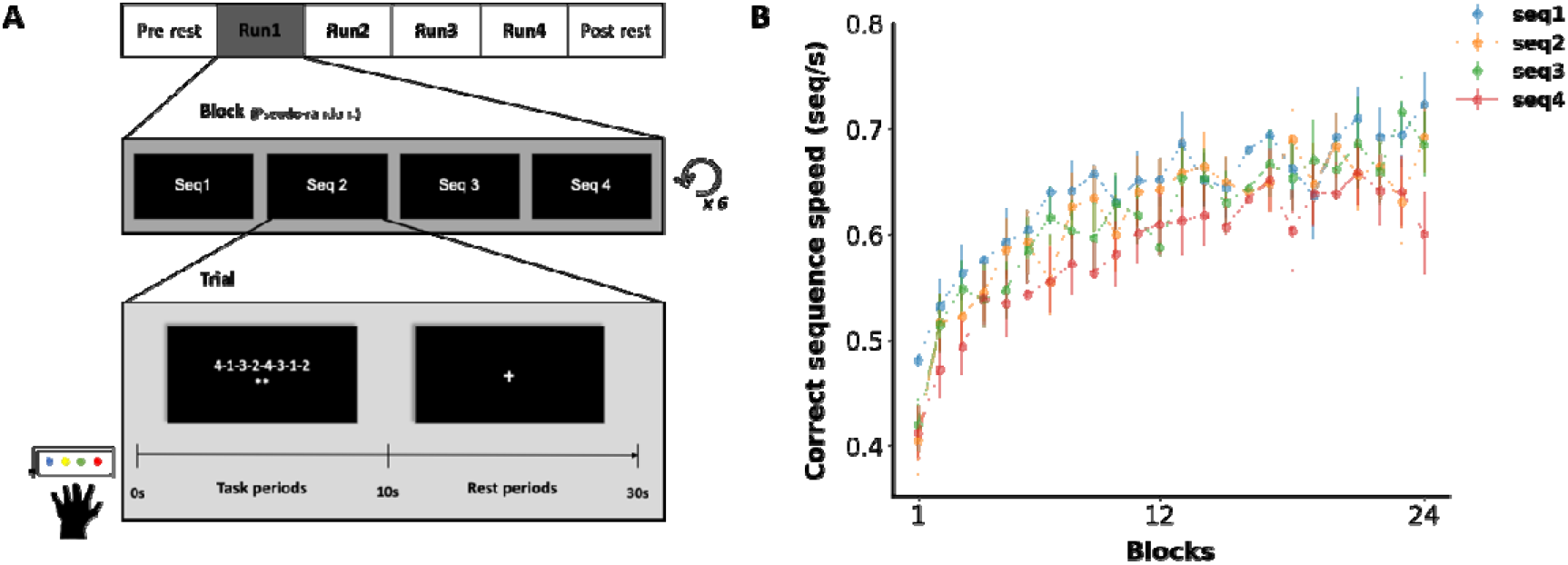
Experiment design and learning curves. (A) Participants conducted a motor sequence learning task in the MRI scanner. Four different finger sequences in each block were presented in a pseudo-random order. Each finger sequence was displayed on the screen during 10 s active task period, which was followed by 20 s rest period. (B) Performance measured as a correct sequence speed improved across multiple task blocks for the four sequences.

Task performance was measured as correct sequence typing speed, calculated as the inverse of the inter-tap interval. All the participants improved their performance for four different sequences, as indicated by faster tapping speeds during the last block compared to the first. There was no significant difference in overall learning performance, which was calculated as an improvement of correct sequence speeds from the first (or second) block to the last block (or second to the last) among the four sequences (*F*(3,72) = 1.43, *P* = 0.24) (Figure 1B, see Methods for details).

### Regions of Interest (ROI) Selection

To identify the regions of interest (ROI) related to the task, we conducted a whole-brain voxel-wise general linear model (GLM) analysis using fMRI data collected from individual participants. We first identified regions where coefficients of GLM regressor encoding the 10 s active task periods (top in Figure 2), which are significantly positive at highly stringent threshold level, *P* < 10^−4^. Then, 14 regions of interest (ROI) were defined as intersections between the significantly positive regions and the predefined sensorimotor regions from a functional atlas^19^. Following previous studies^20,21^, we selected the predefined regions, including bilateral supplementary motor areas (SMA), motor/somatosensory cortices (M1/S1), superior parietal lobules (SPL), and inferior parietal lobules (IPL). The finally defined ROIs were matched to the well-known motor learning network in the brain^20-22^. We used these 14 ROIs in the subsequent fMRI pattern analysis (Figure 2; see more details in Methods).

**Figure 2.**
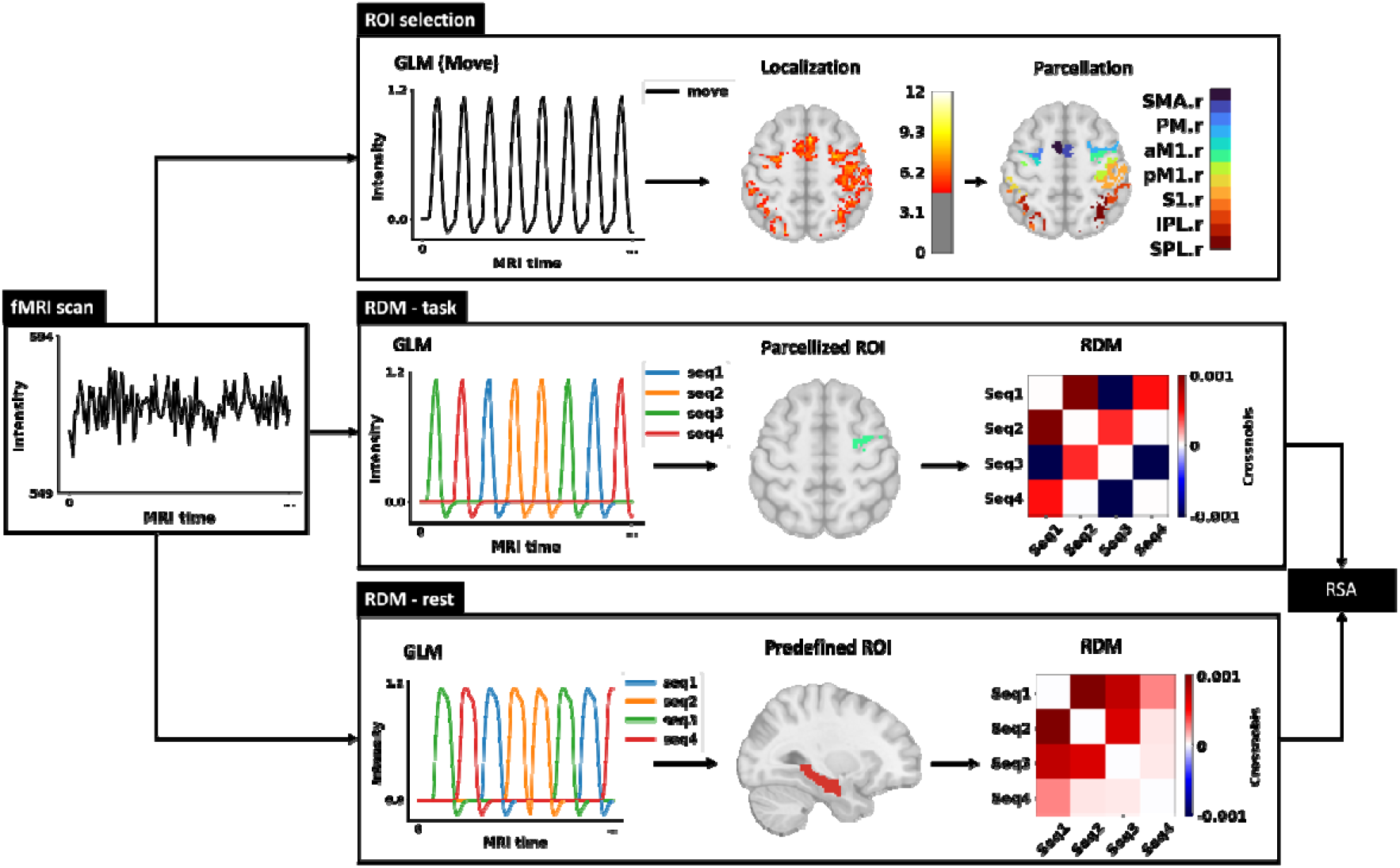
Conceptual diagram of fMRI data analysis. Within the predefined sensorimotor regions from a functional brain atlas, we defined ROIs as regions which were significantly activated during the task using a single GLM regressor encoding active task periods. fMRI activity pattern for each of four sequences was calculated using a separate GLM regressor encoding each sequence’s active task periods. For the defined sensorimotor ROIs, RDMs were reconstructed using crossnobis distance among activity patterns associated with four sequences. Similarly, for the hippocampal ROIs, RDMs were also reconstructed using activity patterns during wakeful rest periods. For the defined ROIs, the pattern separation magnitudes were measured as the mean of lower-triangular elements of RDMs. SMA: Supplementary motor area, PM: Premotor cortex, M1: Primary motor cortex, S1: Primary somatosensory cortex, SPL: Superior parietal lobule, IPL: Inferior parietal lobule, prefix a: anterior, p: posterior.

### Pattern separation in sensorimotor regions

We assessed pattern separation of fMRI activity during the active task periods to investigate how motor sequences were represented in the distinct sensorimotor regions (middle in Figure 2). The pattern separation was measured as the mean fMRI pattern dissimilarity among the motor sequences using a cross-validated squared Mahalanobis distance (i.e., crossnobis distance), known to be unbiased with a zero mean^23^. Specifically, for each of the 14 ROIs, we constructed a representational dissimilarity matrix (RDM), of which element indicates crossnobis distance between the fMRI activities related to a corresponding pair of sequences (Figure 2, see Methods for details). The mean dissimilarity as a measure of pattern separation is the mean value of the lower (or upper) triangular elements of the RDM.

Among the 14 tested ROIs, the mean dissimilarity was the most prominent in the parietal regions, SPLs (left: *T*(24) = 4.42, uncorrected *P* < 10^−4^, corrected *P* = 0.0013, right: *T*(24) = 3.46, uncorrected *P* = 0.001, corrected *P* = 0.014) and IPLs (left: *T*(24) = 3.34, uncorrected *P* = 0.0014, corrected *P* = 0.019; right: *T*(24) = 4.05, *P* < 0.0005, corrected *P* = 0.0032). It was marginally significant in the M1/S1 regions (right S1: *T*(24) = 2.34, uncorrected *P* = 0.014, corrected *P* = 0.18; right anterior M1, aM1.r: *T*(24) = 2.02, uncorrected *P* = 0.027, corrected *P* = 0.32) (Figure 3A). These findings align with the results of a recent study^20^, which suggested that higher-level sequences are not represented in M1/S1, instead in the parietal, SMA, and premotor regions. However, we find significant representation only in the parietal regions, SPLs, and IPLs (Figure 3A). The whole-brain searchlight analysis also identified the parietal regions with the most significant pattern separation, which is consistent with the results of the ROI analysis (Table 1 and Figure 3B). The visual cortex also exhibited significant pattern separation due to distinct visual stimuli of numbers displaying motor sequences. The significant clusters in the dorsolateral prefrontal cortex (DLPFC) would be related to working memory involved in remembering sequences for better performance.

**Table 1.** Clusters with significant pattern separation identified by searchlight analysis.

|  | Peak (MNI) |  |  | Cluster size<br>(voxels) | Z-score<br>at peak | Corrected<br>P-value |
| --- | --- | --- | --- | --- | --- | --- |
|  | X | Y | Z |  |  |  |
| *R Superior Parietal Lobule* | 31 | -63 | 63 | 219 | 4.21 | $P < 0.03$ |
| | 29 | -51 | 43 | 163 | 4.26 | $P < 0.05$ |
| | 41 | -31 | 43 | 129 | 4.30 | $P < 0.08$ |
| *R Dorsolateral Prefrontal Cortex* | 21 | 45 | 49 | 218 | 4.18 | $P < 0.03$ |
| *L Primary Visual Cortex* | -17 | -91 | -9 | 213 | 3.92 | $P < 0.03$ |
| *L Inferior Parietal Lobule* | -35 | -53 | 57 | 204 | 4.50 | $P < 0.03$ |
| *R Inferior Parietal Lobule* | 57 | -43 | 39 | 129 | 4.01 | $P < 0.08$ |

Note: Voxel-wise $P < 0.005$ , 155 suprathreshold voxels as a minimum cluster size for cluster-wise $P < 0.05$

**Figure 3.**
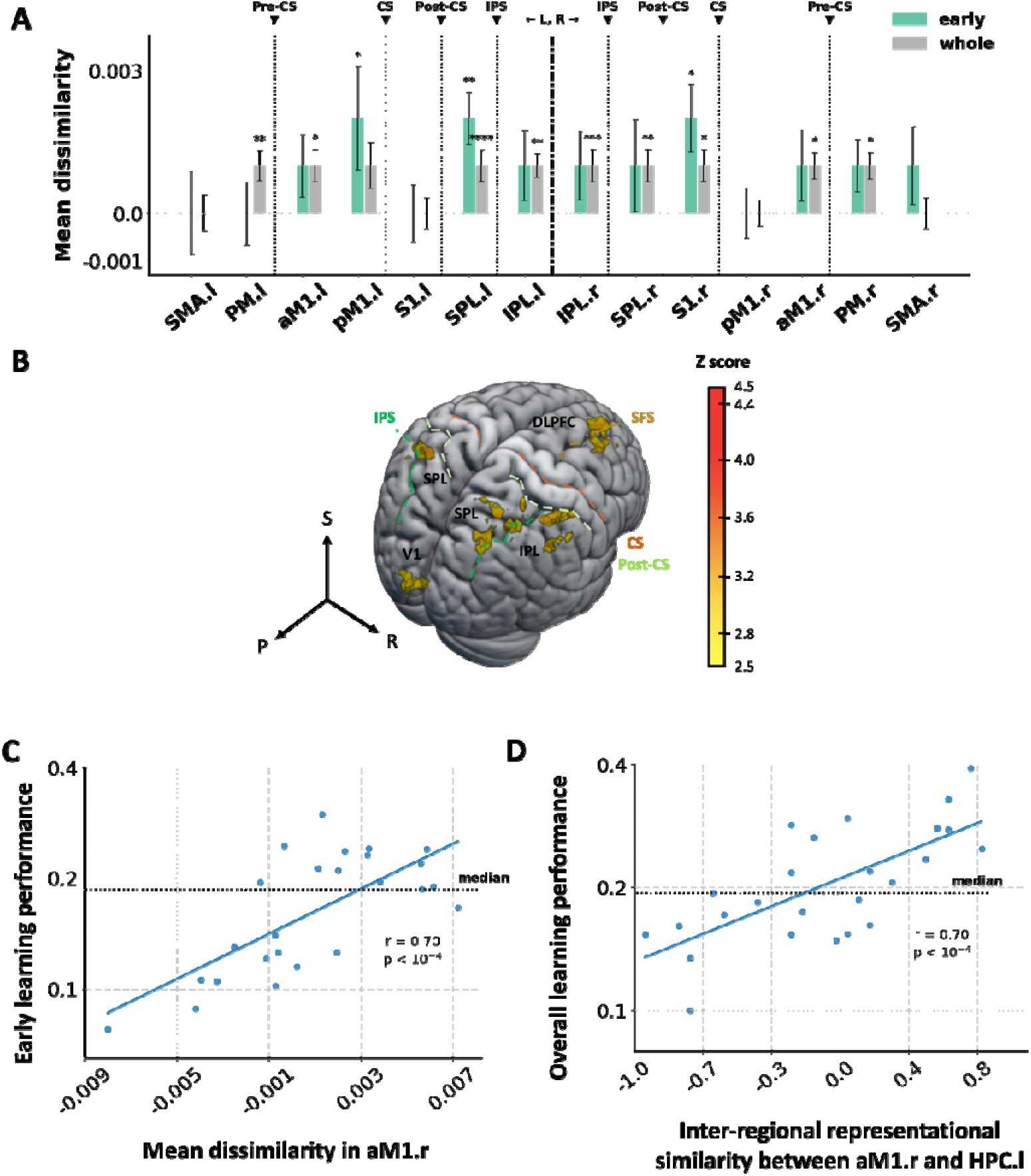
Multivariate fMRI pattern analysis in sensorimotor and hippocampal regions. (A) Mean dissimilarity of fMRI patterns related to motor sequence in the sensorimotor ROIs for early learning and entire learning. (B) Whole-brain searchlight analysis of the mean dissimilarity (see Table 1). (C) Association between early learning performance and pattern separation (i.e., mean dissimilarity) in the right anterior M1 (aM1.r). (D) Association between overall learning performance and aM1.r-HPC.l representational similarity. In (C) and (D), dots indicate individual participants. DLPFC: Dorsolateral prefrontal cortex, V1: Primary visual cortex, CS: Central sulcus, IPS: Intraparietal sulcus, SFS: Superior frontal sulcus, HPC: Hippocampus.

Next, we hypothesized that pattern separation could contribute to learning four distinct motor sequences simultaneously. To test the hypothesis, we correlated mean dissimilarities within the ROIs with individual learning performance (Figure S1), calculated as the difference in tapping speeds for correct sequences between the first and last blocks. Consequently, we found no significant correlation between mean dissimilarities in sensorimotor ROIs and individual learning performance. It suggests that pattern separation primarily contributes to the early stage of learning, where substantial performance improvement occurs. Indeed, the learning curves almost reached a plataeu around the middle of the training blocks (Figure 1B). Intriguingly, the mean dissimilarity in the right (i.e., contralateral) anterior M1 (aM1.r) region highly correlated with individual performance during the early phase of learning (*r* = 0.70, uncorrected *P* < 10^−4^, corrected *P* = 0.0027) (Figure 3C). It was significantly positive for the fast learners (*T*(11) = 3.63, *P* = 0.0020, one-sided) and larger than that for the slow learners (*T*(23) = 2.62, *P* = 0.015, two-sided). The individual early learning performance was highly correlated with the individual overall performance (*r* = 0.74, *P* < 10^−4^), but only the former was significantly related to the pattern separation in the aM1.r.

### Cortico-hippocampal interaction related to the overall learning

Given that pattern separation in the right anterior M1 (aM1.r) during early learning correlated with individual performance, we investigated how the activity patterns in the region were represented in the hippocampus during the rest periods. To this end, we first tested whether the activity patterns related to motor sequences were separately represented in the hippocampus during the rest periods, potentially due to hippocampal replay. However, none of the six hippocampal regions exhibited significant pattern dissimilarity for either early or overall learning (*P* > 0.3). Furthermore, the mean dissimilarities were not correlated with either early or overall learning performance (*P* > 0.05).

Despite these negative outcomes, we alternatively hypothesized that the cortico-hippocampal interaction would still be reflected in representational similarity across the interleaved task and the rest periods. Additionally, we hypothesized that their interaction contributed to learning, as indicated by previous studies demonstrating the contribution of the hippocampus during wakeful rest^10-12^. To test these hypotheses, we first calculated RDMs of the aM1.r during the task periods and the six hippocampal ROIs during the rest periods. Then, for each hippocampal ROI, representational similarity with aM1.r was measured as a second-order similarity between two RDMs using Spearman rank correlation. Then, we correlated the representational similarity with learning performance across 24 participants.

During early learning, the representational similarity between the aM1.r and any of the six hippocampal ROIs was not correlated with individual performance. This finding contrasts with the observation that individual performance was highly correlated with the mean dissimilarity within aM1.r during early learning. However, we discovered that representational similarity between the aM1.r and the entire left hippocampus highly correlated with overall learning performance. (Figure 3D; *r* = 0.70, uncorrected *P* < 10^−4^, corrected *P* = 0.0012). The left hippocampus and aM1.r representational similarity was significant for fast learners (*T*(11) = 3.13, *P* = 0.0048, one-sided) and lager than slower learners (*T*(23) = 4.47, *P* < 0.0005, two-sided). This result further supports the role of M1-hippocampus interaction across alternated task periods and rest periods in performance improvement.

## Discussion

Emerging evidence from recent studies suggests that the hippocampus, long been thought to be independent of motor memories, contributes to motor sequence learning by rapidly consolidating motor memories during wakeful rest. In the current study, we hypothesized that the hippocampal replay of motor sequences, a potential mechanism underlying rapid consolidation, could be reflected in the reactivation of fMRI activity patterns. Based on this idea, we designed an experiment of learning multiple motor sequences simultaneously, distinguishing it from previous studies investigating the role of the hippocampus in motor learning. The experimental design allowed us to analyze activity patterns in sensorimotor regions and those in the hippocampus related to multiple motor sequences and how they correlate with individual learning performance.

We found pattern separation for motor sequences was highly significant in secondary associative regions such as superior and inferior parietal but only marginally significant in the primary somatomotor regions, M1/S1. These results align with a previous study revealing a hierarchical representation of sensorimotor regions, premotor and parietal regions for higher-level information about motor sequences and chunks, and M1/S1 for lower-level information about individual finger movements^20^. Thus, the observed weak pattern separation in M1/S1 may be attributed to the effect of the first finger press, which was different for four sequences in our experiment, not due to the whole sequence itself^24,25^. M1/S1 also do not exhibit learning-related changes in pattern separation^25^. Instead, the representation is shaped by the long-term natural hand use^26^. Intriguingly, however, we found a highly significant association between the pattern dissimilarity and individual learning performance only in the contralateral anterior M1 during the early stage of learning. This result could be due to the role of pattern separation in mitigating contextual interference (CI), which is more pronounced during early learning when individuals are simultaneously learning multiple motor sequences. Pattern separation in the M1 region and its relationship to learning performance pave the way for future studies. It would be fascinating to explore whether professional pianists exhibit greater pattern separation in the M1 region compared to novices^27,28^.

While pattern separation is marginally significant in the anterior M1, it was not significant in the hippocampus during rest periods. However, we found significant representational similarity between the two regions for fast learners and was related to individual overall learning performance. These results indicate an interesting dichotomy for motor sequence representation in M1 and hippocampus. In contrast to our initial hypothesis, the hippocampus did not reactivate distinct motor sequences per se. Instead, it preserved relative differences between sequences represented in the M1 region.

We speculate that the hippocampus reactivates sensorimotor activity patterns in a way that reduces interference among similar motor sequences rather than directly reactivating the motor sequences themselves. Indeed, replay in the hippocampus includes both item-specific and generalizable abstract representation by reorganizing sequences^29^. Due to the task design where multiple sequences were presented in a pseudo-random order, the hippocampus was more likely to be involved in learning abstract and relational structures of sequences instead of representing specific details of sequences^30^. In this way, cortico-hippocampal interaction would reduce interference among sequences or transfer knowledge from one sequence to the other^29-31^, thereby facilitating learning, as observed in our results. This interpretation also aligns with a recent fMRI study demonstrating larger resting functional connectivity between the premotor cortex and the hippocampus under a higher CI practice condition as in our experiment^32^. Interestingly, the cortico-hippocampal interaction in the study was contralateral (left premotor - right hippocampus), which is comparable to our results (right anterior M1-left hippocampus), using the same non-dominant left hand.

Our two main findings of high correlation with individual learning depended on learning stages; pattern separation in the anterior M1 correlated with early learning, and anterior M1-hippocampus interaction correlated with overall learning. Pattern separation in the M1 region would be more critical in early learning since interference among similar motor sequences is initially larger and decreases as learning proceeds. In contrast, the cortico-hippocampal interaction would persist beyond the initial stage of separating sequences, and consolidation occurs throughout the entire learning period. The more sustained interaction between the sensorimotor and hippocampal regions, at first glance, seems to contradict previous studies demonstrating more significant hippocampal replay in early learning where most micro-offline gains occurred. In contrast, in the current experiment, participants learned multiple and longer (8 elements) motor sequences in a pseudo-random order, while the previous studies used a single and shorter (5 elements) sequence. Consequently, we cannot assess the micro-offline gain between two consecutive trials, which the previous studies suggest the effect of hippocampal reactivation during rest periods. Instead, we assessed the learning performance over a longer period and correlated it with cortico-hippocampal interaction. Nonetheless, future studies are warranted to clarify how performance in different learning stages is related to pattern separation and cortico-hippocampal interaction.

The current study has several limitations. First, we used four distinct motor sequences, yielding only six similarity values between sequences, while few previous studies used five fingers (10 values)^26,33^. Although the limited number of sequences might underpower statistical outcomes (e.g., mean dissimilarity) compared to those of a previous study using eight different sequences, the overall pattern separation results were consistent with the previous study^20^. Second, the current experiment design, which has no control group learning sequences in a repeated order, does not allow us to assess interference among sequences directly. Indeed, neural substrates of overnight memory consolidation depend on how multiple motor tasks are presented, either in a random or in a repeated order^5^. A future study would be warranted to investigate how the cortico-hippocampal interaction contributes differently to rapid memory consolidation under high versus low contextual interference. Moreover, there also could be positive effects among sequences, that is, transfer of learning among sequences. In contrast to interference, it could facilitate overall learning, and thus, the association between pattern separation and learning performance should be interpreted with caution.

Taken together, our unique experiment design of learning multiple motor sequences with interleaved rest periods enabled us to reveal how the M1-hippocampus interaction contributed to motor sequence learning. Our results support the idea that reactivation in the hippocampus is not necessarily item-specific. Instead, it is related to learning the abstract relational structure of motor sequences, as evidenced by representational similarity analysis. These findings intrigue future research to explore a novel hypothesis concerning the role of hippocampal replay in multi-task motor learning.

## Methods

### Participants

Twenty-five neurologically healthy and right-handed subjects participated in the study (8 females; ages: mean ± SD = 2.53 ± 3.2, 19 - 35 years). They provided written informed consent and received compensation for their participation. Sungkyunkwan University Institutional Review Board, Suwon, Republic of Korea approved all the procedures of the current study.

### Experiment design

We instructed participants to press four buttons with the fingers of the non-dominant left hand, following a sequence presented on the screen as quickly and accurately as possible. An asterisk indicating a correct press appeared below the sequence of numbers to motivate them to improve performance. We also instructed them to receive as many asterisks as possible. To ensure participants understood the instruction, they performed a behavioral test 1-2-3-4-1 outside the MRI scanner before the experiment. We confirmed that they pressed the buttons following the sequence multiple times and stopped pressing them during the rest periods.

The entire fMRI experiment consists of two resting state runs before and after four main task runs (Figure 1A). We focus on the main task in the current study, and the other resting state data will be reported elsewhere. Each of the main task runs consists of 24 trials; thus, there are 96 trials in total. For each trial, one of four 8-element sequences (1-4-2-3-1-2-4-3, 2-1-3-4-2-3-1-4, 3-4-2-1-3-2-4-1, 4-1-3-2-4-3-1-2, 1 being the pinky finger and 4 being the index finger) was presented for 10 s and followed by a rest period of 20 s with a fixation cross on the center of the screen. We intentionally chose the long inter-stimulus interval (ISI) to ensure enough time for pattern reactivation during the rest. Additionally, this choice would benefit reliable estimation of the activity pattern in the rest period while less affected by a delayed hemodynamic response from the previous task period. Significantly, the slow block design would reduce artifacts in neural representational similarity analysis^34,35^. Four different sequences were presented in a pseudo-random order in a task block, so all four sequences were presented in four consecutive sequences (Figure 1A).

### MRI acquisition

We collected fMRI image data using 3T MRI in the Center for Neuroscience Imaging Research (CNIR) located at Sungkyunkwan University with a 64-channel head coil. An ear plug and a form pad were used to reduce noise from fMRI and participants’ head motion. Functional scans during four runs were acquired by using an echo planar imaging (EPI) sequence with the following spec: repetition time (TR) = 2000 ms; voxel size = 2.0 x 2.0 x 2.0 mm; echo time (TE) = 30 ms; field of view (FOV) = 200 mm; flip angle (FA) = 90º; slice thickness = 2.0 mm; matrix = 96 x 114 x 96 voxels; the number of time steps = 363. Following each of functional run, two EPI images with opposite-phase directions (posterior-to-anterior) were acquired for distortion correction. For anatomical reference, a whole-brain T1-weighted anatomical scan was performed using a magnetization-prepared rapid acquisition with gradient echo MPRAGE sequence, with the following parameters: TR = 2300 ms; voxel size = 1.0 x 1.0 x 1.0 mm; TE = 2.28 ms; FOV = 256 mm; FA = 8° ; slice thickness = 1.00 mm; matrix = 193 x 229 x 193 voxels.

### fMRI data preprocessing

The T1 and fMRI data were preprocessed following a standard procedure suggested by Analysis of Functional NeuroImages (AFNI) software.^36^ The T1 images and functional echo planar image (EPI) data were transformed into volume data using AFNI’s “Dimon” function and realigned from oblique to cardinal orientation using AFNI’s “3dWarp”. The inhomogeneity in the T1 image due to magnetic field bias was corrected, and the skull was removed using AFNI’s “3dUnifize” and “3dSkullStrip”, respectively. Each participant’s brain was transformed into a standard MNI (Montreal Neurological Institute) template to using AFNI’s “@auto_tlrc.” For functional data, spikes were removed using AFNI’s “3dDespike” to reduce outlier effects. Then, time correction was applied to minimize time differences between each slice using AFNI’s “3dTshift”. An affine transformation was used to adjust motion-related artifacts using AFNI’s “3dAllineate”. All the EPI data have a resolution 2.0 x 2.0 x 2.0 mm.

### Sensorimotor and hippocampal ROI selection

Sensorimotor regions of interest (ROI) were extracted from Brainnetome atlas.^19^ We defined 14 ROIs as task-related ROIs: SMA (medial Broadman area 6), PM (entire Broadman area 6 without caudal dorsolateral and ventrolateral of Broadman area 6), aM1(caudal dorsolateral and ventrolateral of Broadman area 6), pM1 (precentral gyrus without caudal dorsolateral and ventrolateral of Broadman area 6), S1 (postcentral gyrus), SPL (superior parietal lobule), and IPL (inferior parietal lobule) from each side of the brain. Within each of the 14 predefined ROIs, we further defined subregions related to the motor sequence task using a generalized linear model (GLM) analysis. A design matrix of GLM included a boxcar regressor encoding task periods (“Move” regressor) for all sequences, regressors of non-interest modeling six head motions, and third-order polynomials of fMRI signal drift. We performed the first-level GLM analysis for each of the four fMRI runs and 25 participants. Then, four activation maps of the “Move” regressors were averaged for each participant and then entered for the second-level group analysis using AFNI’s 3dttest++ function. We used highly stringent voxel-wise p-value < 10^−4^ for the significance level to define the task-related ROIs.

The hippocampal ROIs included rHPC (rostral hippocampus) and cHPC (caudal hippocampus) on each brain side. These ROIs were also extracted from the Brainnetome atlas. These ROIs were chosen based on our hypothesis that the hippocampus would contribute to motor learning during rest periods.

### Activation maps for pattern analysis

To calculate the activation level of each sequence, a separate GLM analysis was performed for each task and rest periods. Having two regressors encoding both periods in a single GLM could cause a multicollinearity problem due to the overlap of regressors convolved with the hemodynamic response. GLM analyses were conducted separately on the task and rest periods to address this issue. The GLM for the task periods included four regressors encoding different sequences separately and regressors of non-interest, including six regressors related to head motions and third-order polynomials modeling fMRI signal drifts. The GLM for the task periods was designed similarly, except for four regressors encoding different rest periods following sequences. GLM analyses were performed separately for each of the four fMRI runs, producing eight activation maps for four task periods and four rest periods following sequences.

### Representational pattern analysis

To construct a representational dissimilarity matrix (RDM), we computed a multivariate distance between a pair of activation maps related to four sequences during task periods or resting periods in a specific region of interest. For a distance metric, we selected a cross-validated Mahalanobis distance, which has been shown to be unbiased and reliable due to cross-validation and multivariate noise normalization^23^. The multivariate noise normalization was performed by estimating the covariance from the GLM fitting residuals and regularizing it through Ledoit-Wolf optimal shrinkage^37^Pattern separation, which indicates separability between fMRI activities related to sequences, was quantified as the mean dissimilarity among sequences (i.e., mean of the lower or upper triangular elements of RDM). We assessed the statistical significance of pattern separations by conducting a one-sided T-test across 25 participants to determine whether the mean dissimilarity is significantly larger than zero^38^. We also corrected significance using Bonferroni-correction for multiple tests for 14 sensorimotor ROIs and six hippocampal ROIs.

For searchlight analysis of pattern separation, we used spherical ROIs spanning the whole brain^39^ with a radius of 6 mm (93 nearest voxels from a center, 744 mm^2^), all within the brain mask. Likewise, in the ROI analysis, we conducted a one-sided T-test for mean dissimilarities across participants for searchlight ROIs in the whole brain. Monte Carlo simulation using a voxel-wise threshold *P* < 0.005 determined a minimum cluster size of 155 suprathreshold voxels (1240 mm^2^) connected through faces, edges, or corners for cluster-wise corrected threshold *P* < 0.05 within the whole-brain gray matter mask. We used AFNI’s “3dttest++” with “-Clustsim” option for this cluster analysis (Table 1).

For the inter-regional representational similarity analysis (RSA), we calculated second-order similarity using Spearman correlation between the RDMs of the right anterior motor cortex (aM1.r) during task periods and each RDM of the six hippocampal ROIs during rest periods (Figure 2). The significance level is calculated by a one-sided T-test whether the correlation is larger than zero. Pattern separation (i.e., mean dissimilarity) and aM1.r-hippocampal representational similarity were correlated with early learning and overall learning performance. For all the ROI analyses, we used Bonferroni-correction for multiple tested ROIs and the two different learning performances. Specifically, there were 28 and 12 multiple tests, respectively for sensorimotor ROIs and hippocampal ROIs.

### Individual learning performance

For each sequence, performance (i.e., correct sequence speed) was measured as the number of correct sequences per second, which was the inverse of time spent to complete an 8-element sequence correctly. We averaged the correct sequence speed for four sequences presented in a block. If there was no successfully tapped trial for any sequence, we excluded the sequence when averaging. The early learning performance was measured as a difference between the correct sequence speeds in the first block of the first fMRI run and the last block of the second fMRI run. For the overall learning, we used the last block of the last (fourth) fMRI run instead of the second. To compare the overall performance among sequences, we used the second block or the second to the last block if there was no successful trial for at least one sequence in the first or the last block. We defined 12 fast learners whose performance is larger than the median performance among 25 participants.

## Data and code availability

All MRI data used in this study were archived in the Hanyang University Network Attached Storage (NAS). If necessary, the corresponding author can provide any information on the dataset and codes related to this study.

## Acknowledgments

This work was supported by the National Research Foundation of Korea (NRF-2021R1A2C2011648), Hanyang University (HY-202000000002753), and Center for Neuroscience Imaging Research, Institute for Basic Science, Korea (IBS-R015-D1). Neuroimaging was performed at the Center for Neuroscience Imaging Research located at Sungkyunkwan University, supported by Institute for Basic Science. We thank Antoine Caraballo for his helpful comments and discussion.

## Author contributions

S.K. conceived of the idea presented in the study. S.Y. and S.K. designed the experiment. S.Y. carried out the experiment and formal analysis; S.Y. and S.K. wrote the manuscript. During the preparation of this work, the authors used ChatGPT4.0 and Grammarly in order to edit the manuscript with grammar checks. After using these tools, the authors reviewed and edited the content as needed and take full responsibility for the content of the publication.

