## Supplementary material for "Cortico-hippocampal representational similarity contributes to learning of multiple motor sequences": Figure S1

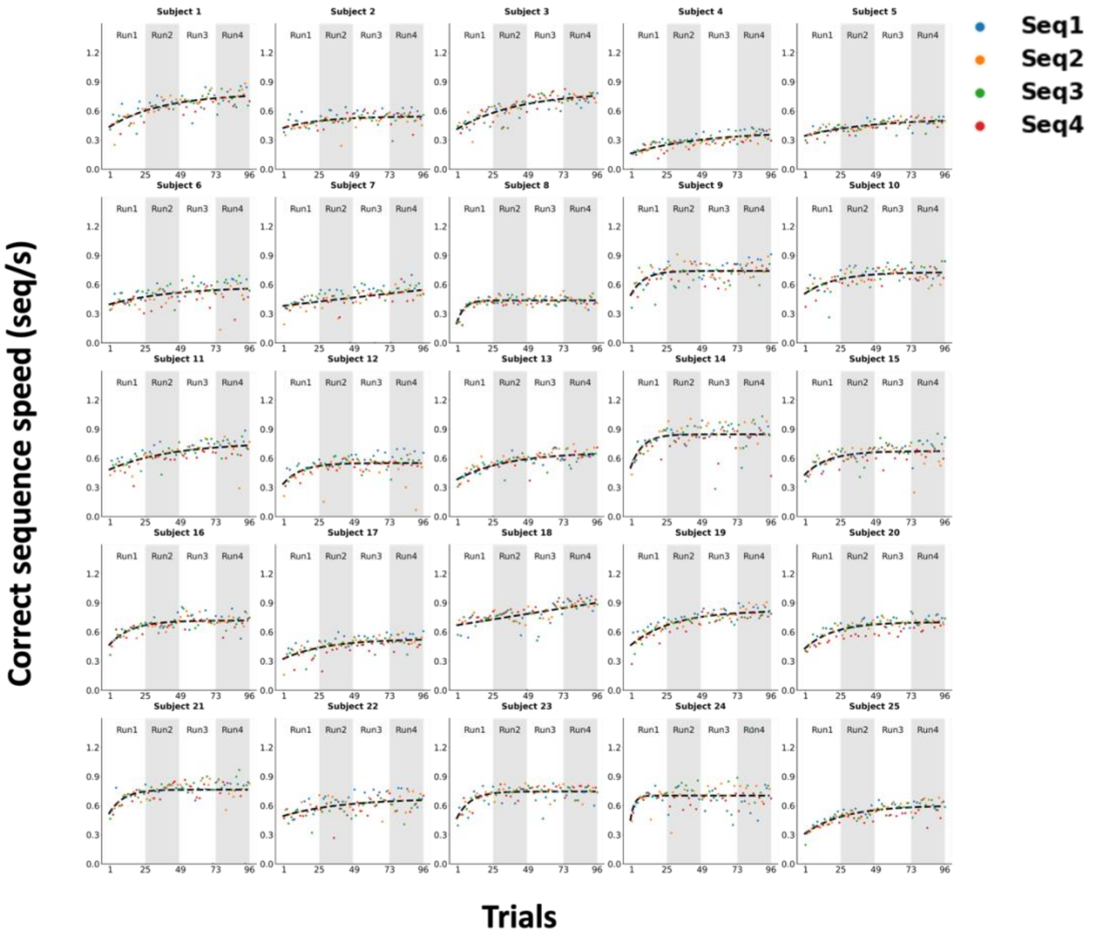


**Figure S1. Learning curves of 25 individual participants**

For all the 25 participants, performance measured as a correct sequence speed improved across multiple task blocks for the four sequences. We correlated pattern separation in the regions of interest with the individual early and overall learning performance.
